# Frizzled-dependent Planar Cell Polarity without Wnt Ligands

**DOI:** 10.1101/2020.05.23.108977

**Authors:** Joyce J.S. Yu, Aude Maugarny-Calès, Stéphane Pelletier, Cyrille Alexandre, Yohanns Bellaiche, Jean-Paul Vincent, Ian J. McGough

**Affiliations:** The Francis Crick Institute, London NW1 1AT, UK; Institut Curie, PSL Research University, CNRS UMR 3215, INSERM U934, F-75248 Paris Cedex 05, France; Sorbonne Universités, UPMC Univ Paris 06, CNRS, UMR 3215, INSERM U934, F-75005 Paris, France

**Keywords:** Wnt, Frizzled, PCP

## Abstract

Planar cell polarity (PCP) organizes the orientation of cellular protrusions and migratory activity within the tissue plane. PCP establishment involves the subcellular polarization of core PCP components. It has been suggested Wnt gradients could provide a global cue that coordinates local PCP with tissue axes. Here we dissect the role of Wnt ligands in the orientation of hairs of Drosophila wings, an established system for study of PCP. We found that PCP was normal in quintuple mutant wings that rely solely on membrane-tethered Wingless for Wnt signaling, suggesting that a Wnt gradient is not required. We then used a nanobody-based approach to trap Wntless in the endoplasmic reticulum, and hence prevent all Wnt secretion, specifically during the period of PCP establishment. PCP was still established. We conclude that, even though Wnt ligands could contribute to PCP, they are not essential, and another global cue must exist for tissue-wide polarization.

## Introduction

Planar cell polarity (PCP) refers to the polarity that epithelial cells acquire along the plane of the epithelium, orthogonal to the apical-basal axis. In a wide range of metazoans, PCP contributes to cell proliferation, cell fate decisions, body axis elongation and morphogenesis, as well as to the orientation of cellular protrusions such as hairs (Goodrich and Strutt, 2011; Adler, 2012; Yang and Mlodzik, 2015; Butler and Wallingford, 2017; Lawrence and Casal, 2013). Moreover, aberrant PCP has been linked to disease such as polycystic kidney disease, deafness, and cancer (Yates *et al*., 2010; Lu and Sipe, 2016; VanderVorst *et al*., 2018). Understanding the molecular basis of PCP establishment and maintenance is therefore important from both a fundamental and a biomedical perspective. Genetic analyses have identified many conserved molecules that mediate PCP from flies to humans. A number of these proteins make up the so-called core PCP pathway. They include, for example, the transmembrane proteins Frizzled (Fz1, FZD in vertebrates) and Starry Night (Stan aka Flamingo, CELSR in vertebrates) and the cytoplasmic protein Dishevelled (Dsh, DVL in vertebrates). A second PCP pathway, named after two of its components Fat (Ft) and Dachsous (Ds), has been identified in Drosophila (Mahoney *et al*., 1991; Clark *et al*., 1995; Zeidler, Perrimon and Strutt, 1999). Its involvement in vertebrates is less well characterised than in flies and its relationship with the core pathway remains controversial (Thomas and Strutt, 2012; Matis and Axelrod, 2013). However, the finding that, in some tissues, this pathway can drive PCP in the absence of the core pathway (Casal, Lawrence and Struhl, 2006) shows that cells can use multiple inputs to orient themselves.

Core PCP relies on the complementary localisation of components at the distal and proximal sides of cells. These components include transmembrane proteins, which mediate the cell interactions that coordinate polarity locally, and intracellular factors, which stabilise the asymmetry within each cell (Vinson, Conover and Adler, 1989; Klingensmith, Nusse and Perrimon, 1994; Taylor *et al*., 1998; Wolff and Rubin, 1998; Usui *et al*., 1999; Chae *et al*., 1999; Wallingford *et al*., 2000; Feiguin *et al*., 2001; Strutt, 2001; Axelrod, 2001; Tree *et al*., 2002; Jenny *et al*., 2003; Bastock, Strutt and Strutt, 2003; Jenny *et al*., 2005; Devenport and Fuchs, 2008; Strutt, Warrington and Strutt, 2011). In addition, it is thought that tissue-wide global cues provide an overall direction to PCP relative to embryonic axes. One attractive possibility is that this is achieved by morphogen gradients, although this has not been directly demonstrated. Among the key components of core PCP are Frizzled proteins, transmembrane proteins that bind Wnts through their extracellular cysteine rich domain (CRD) (Bhanot *et al*., 1996; Hsieh *et al*., 1999). Indeed, Frizzled receptors are key mediators of canonical Wnt signalling. Thus, it is conceivable that Frizzled proteins could read and transduce a Wnt gradient into PCP. It has been suggested therefore that a long range Wnt gradient would bias Frizzled activity and kickstart the molecular interactions that stabilise the asymmetric distribution of other core PCP components and the morphological implementation of PCP (Figure 1A) (Gubb and García-Bellido, 1982; Adler, Krasnow and Liu, 1997; Strutt, 2001; Struhl, Casal and Lawrence, 2012; Fisher and Strutt, 2019).

**Fig. 1.**
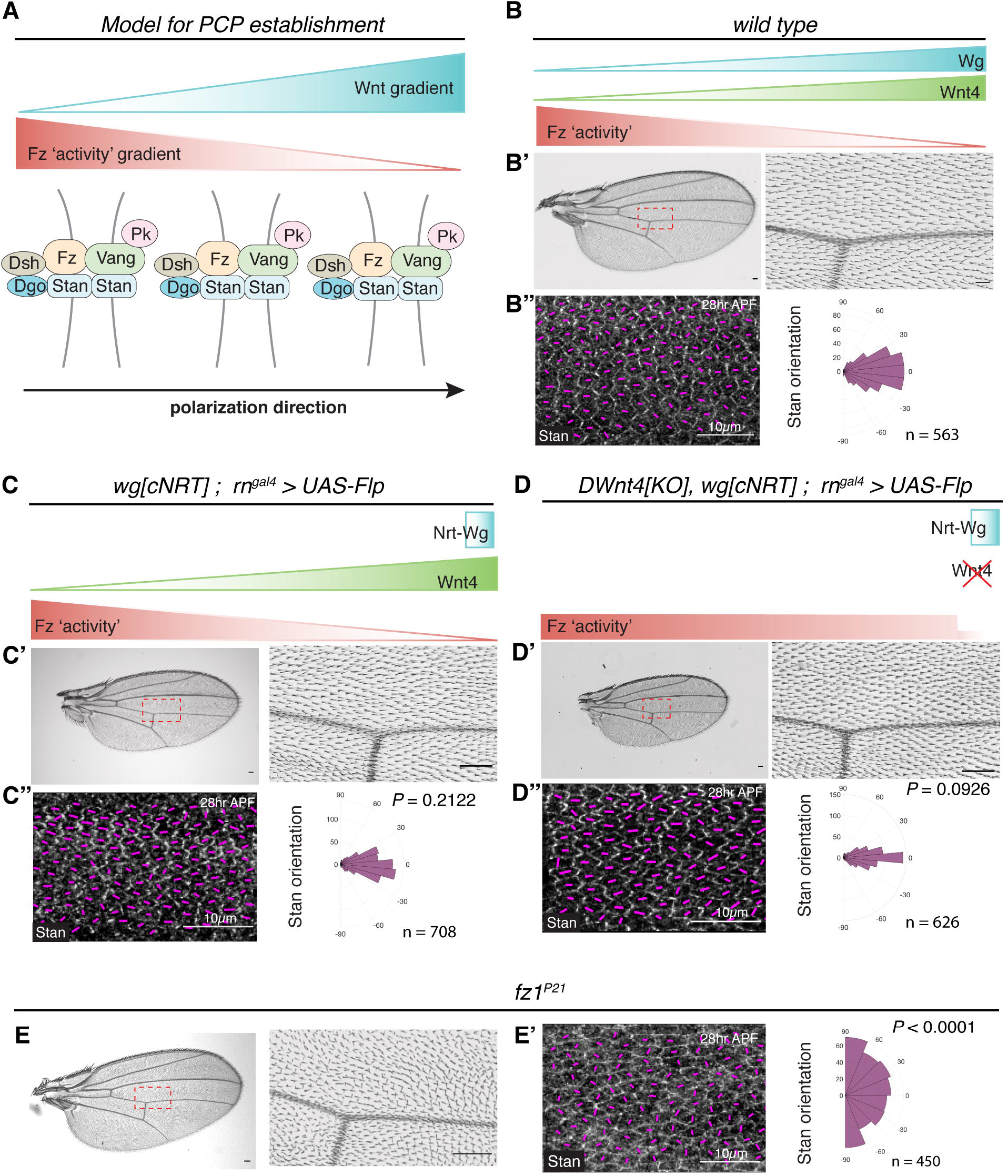
PCP is normal with membrane-tethered Wingless and the absence of DWnt4. (**A**) A model of how a Wnt gradient may lead to a Fz ‘activity’ gradient that directs the asymmetric localisation of core PCP protein complexes, eventually polarising hair outgrowth in one direction. (**B**) A wild type adult wing (**B’**) and a 28hr APF pupal wing with anti-Stan staining (**B’**’). Here and in subsequent figures, the orientation and magnitude of the asymmetric localization of Stan in each cell is depicted by magenta lines. Stan orientation data from the pupal wing region (corresponding to the region marked by a red rectangle in the adult wing) was tabulated on a polar coordinate histogram, with n denoting the number of cells (pooled from 4 pupal wings). (**C, D**) Wing, pupal wing, and Stan orientation polar histogram for homozygous NRT-Wg larvae (no diffusible Wingless) (**C**) or for homozygous NRT-Wg larvae lacking DWnt4 (**D**). P values were calculated with a two-sample Kolmogorov-Smirnov test that compares the distribution of the Stan orientation in mutant conditions (**C”** and **D”**) with that in the wild type (shown in **B’’**). (**E**) Wing from a hemizygous *fz1^P21^* mutant fly. (**F**) Pupal wing and polar histogram of Stan orientations for the same genotype. Scale bars are 50μm unless specified otherwise.

A number of studies have implicated Wnt ligands in PCP establishment. In fish and frog embryos, mutation or knock down of Wnt5a or Wnt11 lead to a reduction in the PCP-dependent cell movements that lead to axis elongation (Rauch *et al*., 1997; Heisenberg *et al*., 2000; Tada and Smith, 2000; Wallingford, Vogeli and Harland, 2001; Andre *et al*., 2015). Likewise, in the mouse limb bud, deletion of Wnt5a interferes with the establishment of PCP in chondrocytes along the proximal-distal axis and thus with tissue elongation (Gao *et al*., 2011; Yang *et al*., 2017). Therefore, in these instances, a Wnt ligand is required for PCP, although it is not clear whether this role is instructive or permissive (Gao, 2012; Lawrence and Casal, 2013). In fish and frog embryos, the PCP phenotypes caused by Wnt inhibition can be rescued by uniform expression of exogenous Wnts, suggesting that graded ligand distribution may not be required for PCP. Nevertheless, localised ectopic localised Wnt expression can orient PCP both in Xenopus mesoderm and the mouse limb bud (Chu and Sokol, 2016; Minegishi *et al*., 2017). Therefore, in these systems, a Wnt gradient can provide information for polarization, even though it may not be required.

Because of a wealth of genetic tools and a long history of PCP research, Drosophila is well suited to investigate the requirement of Wnt gradients in PCP. In Drosophila, PCP is readily assayed by measuring the orientation of hairs that decorate much of the cuticle. For example, at the surface of developing wings, the core PCP pathway orients actin protrusions, which serve as a template for the formation of hairs during metamorphosis (Wong and Adler, 1993). In the wild type, these protrusions point distally but can be reoriented by ectopic expression of Wingless or DWnt4 (Wu *et al*., 2013). Therefore, as in vertebrates, gain-of-function evidence suggests that a Wnt gradient is sufficient to reorient hairs. Loss-of-function tests are difficult to perform and interpret because of the requirement of Wingless for wing specification and growth, which occur before PCP establishment (Ng *et al*., 1996). Nevertheless, it has been possible to interfere with *wingless* and DWnt4 with a combination of hypomorphic alleles compatible with growth. Adult flies were rarely recovered, but in most pupal wings, the pre-hair actin bundles were found to be partially misoriented (Wu *et al*., 2013). This observation suggests that, in the developing wing, Wingless and DWnt4 could be needed redundantly for the establishment of PCP, although it does not directly address whether these Wnts must be graded. In another tissue, the adult abdomen, these two ligands appear to be entirely dispensable for bristle orientation (Lawrence, Casal and Struhl, 2002; Chen *et al*., 2008). Therefore, the requirement of Wnts, and/or the instructive value of their graded distribution for PCP remains a matter of debate both in flies and vertebrates.

The Drosophila genome encodes seven Wnts. Six of them carry a palmitoleate moiety that is essential for engagement with the Frizzled CRD and have a well-defined vertebrate ortholog (in parenthesis below): Wingless (Wnt1), DWnt2 (Wnt7), DWnt4 (Wnt9), DWnt 5 (Wnt5), DWnt6 (Wnt6), and DWnt10 (Wnt10) (Nusse, 1997-2020). Drosophila also encodes a non-conserved WntD, which is not palmitoleoylated and therefore unlikely to bind the CRD of Fz (Wu and Nusse, 2002). We took advantage of recent developments in genome engineering to create a panel of alleles, some conditional, in all the genes encoding Drosophila Wnts. With these genetic tools, we found that PCP can be established in the absence of a diffusion-based Wnt gradient. In further analysis, we show that the Frizzled-dependent core PCP pathway does not need any Wnt ligand to organise PCP in the developing wing. Therefore, in this instance, another non-Wnt global cue must control the subcellular asymmetry of Fz and other core PCP components.

## Results

### DWnt4 is not required for PCP, even in the absence of diffusible Wingless

Larvae expressing membrane-tethered (non-diffusible) Wingless (NRT-Wg) from the endogenous locus can develop into flies with apparently normal appendages, albeit with a delay and at a reduced frequency (Alexandre, Baena-Lopez and Vincent, 2014). Wing hairs in these animals appear to be normally oriented, suggesting that PCP establishment does not require a Wingless gradient. To improve viability and sample recovery, we took advantage of an allele, here referred to as *wg[cNRT]*, which can be converted in a tissue-specific manner from expressing wildtype Wg to expressing NRT-Wg. Allele conversion was induced with *UAS-Flp* and *rn^gal4^*, which is expressed specifically in wing primordia at the 2^nd^ instar stage (St Pierre *et al*., 2002). The resulting pupal wings, which presumably lack diffusible Wingless, were stained for Stan (aka Fmi), an atypical cadherin that forms homophilic bridges across the proximal-distal cell junctions and thus serves as an early marker of PCP (Lu *et al*., 1999; Usui *et al*., 1999). The distribution of Stan was indistinguishable from that in wild type larvae (Fig 1B-C), confirming that diffusible Wingless is not necessary for PCP. It has been suggested that a DWnt4 gradient could redundantly promote PCP in the Drosophila wing (Wu *et al*., 2013). To probe this suggestion, we generated a *DWnt4* mutant (*DWnt4[KO]*) in the *wg[cNRT]* background (Figure S1A). The resulting flies were used to create NRT-Wg-expressing wing primordia lacking all DWnt4 protein. PCP, as determined with Stan localisation, was still normal (Figure 1D). For comparison, significant polarity defects were seen in *fz1^P21^* mutants, as expected since Fz1 is the only Frizzled receptor required for PCP in Drosophila (Figure 1E, F). We conclude that gradients of diffusible Wingless and DWnt4 are dispensable for PCP, but we cannot exclude the possible contribution of other DWnts.

### Multiple Wnts are expressed in the developing wing

To identify the Wnts that are expressed in wing primordia and hence possibly involved in PCP in this tissue, we generated a panel of reporter genes. CRISPR-Cas9 was used to insert DNA fragments encoding nuclear targeted GFP or GAL4 at the endogenous translation initiation codon of DWnt2, DWn4, DWn5, DWn6, DWn10, and WntD, thus allowing transcriptional activity to be readily assessed. DNA encoding an HA-tag was also inserted in the coding region of DWnt10 to generate a protein reporter (Figures S1B-D). Analysis of late third instar wing imaginal discs showed that DWnt4 and DWnt6 are expressed, like Wingless, at the dorsal-ventral (DV) boundary (Figure 2A). DWnt2 was similarly expressed but more broadly. Expression of HA-DWnt10 was undetectable by anti-HA immunofluorescence. However, a weak GFP signal was produced in *DWnt10[GAL4] UAS-GFP* wing primordia, indicating low level expression (Figure S1E). The *DWnt5* and *WntD* reporters remained silent in wing primordia (Figure 1B and S1F). Since these Wnts do not bind the Fz CRD (Wu and Nusse, 2002), we conclude that they are unlikely to contribute to activation of the core Fz PCP pathway. In contrast, the genes expressed in imaginal discs (DWnt2, DWnt4, DWnt6 and DWnt10, and Wingless) continued to be expressed at pupal stages (Figure 1B, S1E), suggesting that any of them could play a role in PCP beyond the period of patterning and growth.

**Fig. 2.**
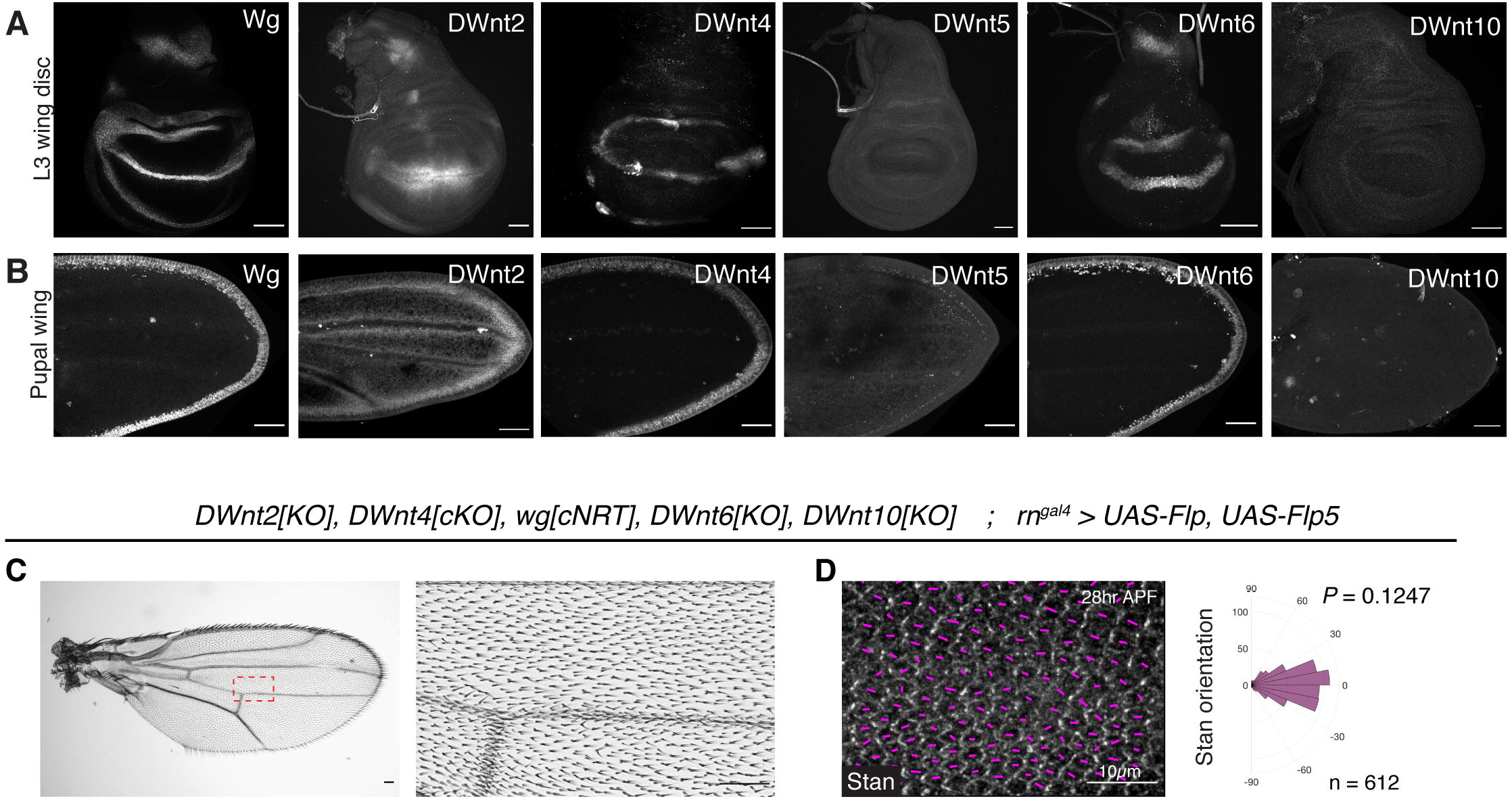
Deletion of all expressed DWnts does not impair PCP in the presence of membrane-tethered Wingless. (**A, B**) Expression patterns of DWnts in third instar wing discs and pupal wings. (**C**) Wing derived from primordia expressing NRT-Wg instead of Wg, and lacking DWnt2, DWnt4, DWnt6 and DWnt10. (**D**) Pupal wings and polar histogram of Stan orientations for the same genotype. The P value was calculated using the two-sample Kolmogorov-Smirnov test to compare the distribution of Stan orientation of the mutant condition (**D**) to that of wild type in Figure **1B’’**.

### PCP establishment in the absence of a Wnt gradient

To test if any Wnt gradient contributes to PCP in the developing wing, we sought to abrogate the activity of *DWnt2*, *DWnt4*, *DWnt6* and *DWnt 10* in the *wg[cNRT]* background. All these genes, except for *DWnt2*, are located within 100kb of each other in the genome, excluding the possibility of recombining individual mutants (Figure S2A). We opted therefore for iterative rounds of CRISPR-Cas9-mediated gene targeting to generate multiple *DWnt* mutants in the *wg[cNRT]* background. First, we sequentially introduced indels at *DWnt6* and *DWnt10* on a chromosome carrying *wg[cNRT]*. Then we generated a conditional *DWnt4* allele (DWnt4[cKO]) in the background of *wg[cNRT], DWnt6[KO], DWnt10[KO]* to generate a quadruple mutant chromosome (*Dwnt4[cKO], wg[cNRT], DWnt6[KO], DWnt10[KO]*). A deletion of the first exon of *DWnt2*, which is located elsewhere on chromosome 2, was generated separately and introduced on this chromosome by standard recombination to generate a quintuple mutant chromosome. Thus, with *rn^gal4^*-driven Flp expression, we obtained wing primordia expressing NRT-Wg instead of Wg and lacking DWnt2, DWnt4, DWnt6 and DWnt10, effectively removing all diffusible Wnts. The resulting fly wings were smaller than wild type (Figure 2C), as previously reported for NRT-Wg wings (Alexandre, Baena-Lopez and Vincent, 2014) but, remarkably, wing hair orientation and the distribution of Stan were normal (Figure 2D). This finding suggests that a Wnt diffusion gradient is not necessary for the establishment and maintenance of Fz-dependent PCP.

### PCP without secreted Wnt

In light of the previous result, we wondered whether Wnts (graded or otherwise) are at all required for the establishment of PCP in Drosophila wings. Since Wingless is required for wing specification and growth, complete and early removal of all Wnts leads to the absence of wing primordia, precluding an assessment of PCP. However, the core Fz PCP pathway is thought to be required only from early pupal stages, after most growth has taken place. Indeed, sensitive imaging techniques have shown that PCP domains start aligning along the proximal-distal axis from late third instar larval stage (Sagner *et al*., 2012; Aigouy *et al*., 2010). Moreover, the PCP phenotype of *fz* mutant could be rescued by uniform Fz-GFP expression up untill 6hr after prepupae formation (APF) (Strutt and Strutt, 2002). Therefore, it appears that the roles of Wnt signalling in growth and PCP can be temporally separated. We tested this further with a conditional allele of *dsh*, which is required for both activities (Figure S3A, B). Inactivation of this allele (*dsh[cKO]*) with *UAS-Flp* and *nub^gal4^*, which is expressed specifically in wing primordia at the late second instar stage (Zirin and Mann, 2007) allowed sufficient growth to reveal the expected PCP phenotype (Figure S3C-E). There is therefore a temporal window when the role of Wnt ligands in PCP can be assessed independently of their role in growth.

All Wnts (except WntD) require the multi-pass transmembrane protein Wntless/Evenness-interrupted (here referred to as Wls) for progression in the secretory pathway (Banziger *et al*., 2006; Bartscherer *et al*., 2006; Herr and Basler, 2012). Complete loss of Wls effectively prevents the secretion of all Wnts and experimental abrogation of Wls could therefore be used to inhibit the activity of all Wnts at once. However, Wls protein activity is known to perdure (Banziger *et al*., 2006; Bartscherer *et al*., 2006), limiting the temporal resolution of a conditional allele or RNAi-mediated interference. To overcome this limitation, we designed an approach to target the Wls protein as well as the gene. Inhibition of Wls protein was achieved by trapping it in the endoplasmic reticulum (ER), thereby preventing its progression, and that of all Wnts, through the secretory pathway. We first engineered the *wls* locus so that it expressed a functional GFP fusion, with the GFP moiety on the luminal side (*wls[ExGFP]*) (Figure S3F). We also created a transgene for Gal4-dependent expression of an anti-GFP nanobody modified to be retained in the ER lumen (*UAS-Nanobody^KDEL^*) (Figure 3A). Homozygous *wls[ExGFP]* larvae expressing this transgene under the control of *vg^gal4^*, an early wing primordium driver, gave rise to flies lacking wings (Figure 3B), the same phenotype seen in *wingless^1^* mutants, which lack the wing enhancer of *wg* (Sharma and Chopra, 1976). Hence, trapping Wls in the ER is an effective approach to inhibit Wnt secretion.

**Fig 3.**
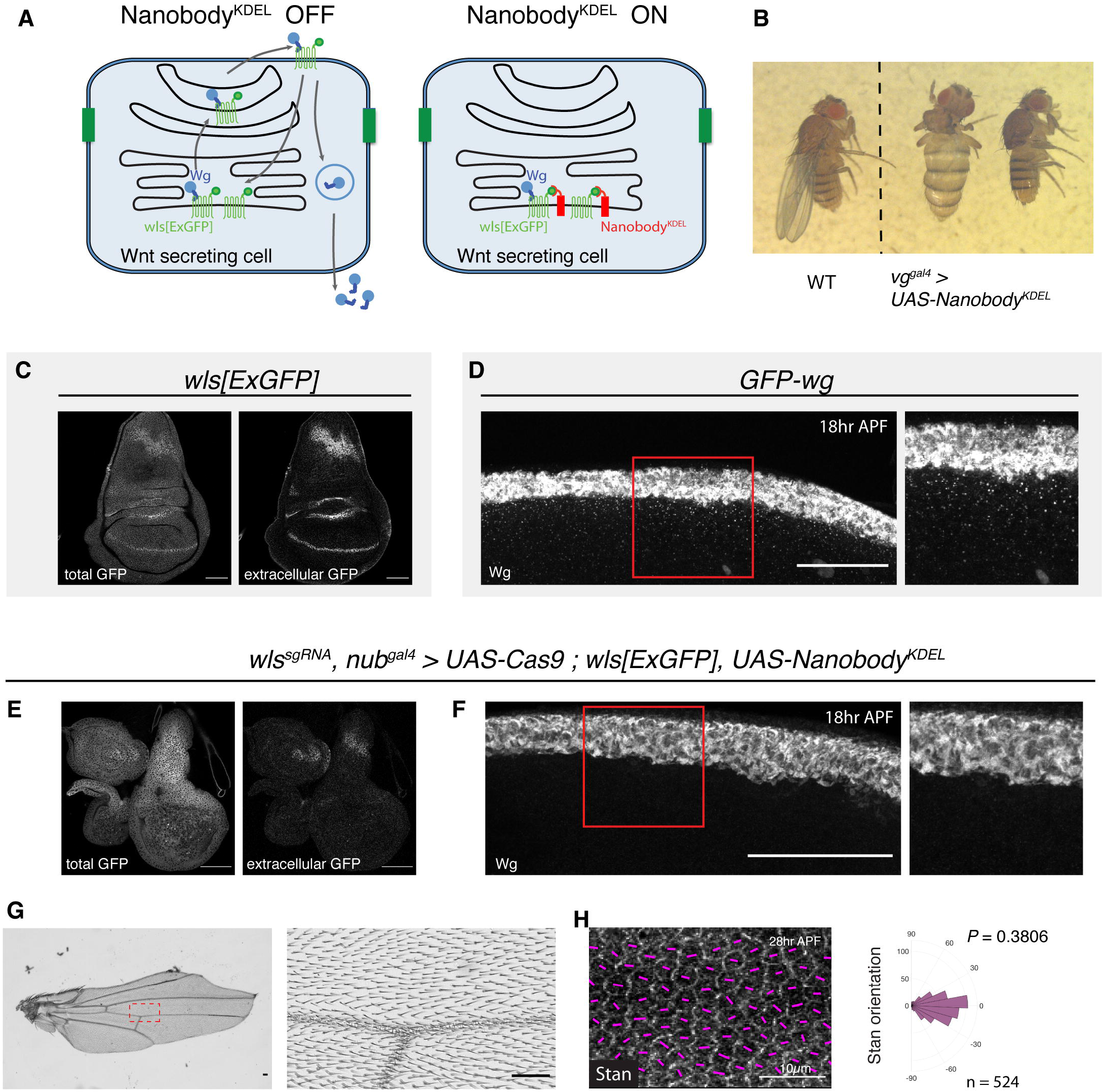
Inactivation of all DWnts during the mid-third instar does not impair PCP. (**A**) Diagram showing how Nanobody^KDEL^ is expected to prevent Wnt secretion. (**B**) Expression of Nanobody^KDEL^ specifically in wing primordia and from the onset of development (with vg^gal4^) in homozygous *wls[ExGFP]* larvae phenocopies a *wingless* mutant. (**C**) Total and extracellular Wls[ExGFP] in a third instar *wls[ExGFP]* wing disc. This recapitulate the pattern seen with wild type Wls. (**D**) GFP-Wg 18hr APF pupal wing showing Wingless diffusion away from the wing margin. (**C-H**) All DWnts were inactivated by driving *Nanobody^KDEL^* expression and *UAS-Cas9* with the wing pouch driver *nub^gal4^*. The driver, which is located on the 2^nd^ chromosome was used instead of the previously used rn^gal4^ (3^rd^ chromosome) to overcome constraints caused by the number of transgenic alleles already present on the 3^rd^ chromosome. (**E**) Wing disc showing the relative absence of Wls within the pouch (compared to wild type in **C)**. (**F**) immunoreactivity in an 18hr APF pupal wing, showing the lack of signal away from the wing margin (compare to **D)**. (**G, H**) Adult wing and Stan polarization in pupal wings when all DWnts were inactivated.

Having established the effectiveness of the Wls trapping approach, we turned to the nub^gal4^ driver to activate *UAS-Nanobody^KDEL^* after sufficient growth has taken place. This was combined with a tissue-specific gene knockout technique involving expression of *UAS-Cas9* in the presence of a transgene expressing a guide RNA targeting *wls* (Port and Bullock, 2016; Port *et al*., 2020). Thus, in the *wls[ExGFP]* background, one Gal4 driver suffices to induce expression of Cas9 (for gene inactivation) and Nanobody^KDEL^ (for sequestration of the protein product). In the resulting imaginal discs, Wls-GFP was no longer detectable at the cell surface (compare Figure 3C and E), confirming its depletion/ trapping. Moreover, in the resulting pupal wing, Wingless was retained within Wingless-producing cells (compare Figure 3D and F) and therefore unable to activate signal transduction. Indeed, adult wings of this genotype lacked margin tissue (Figure 3G), which is specified by canonical signalling (Couso, Bishop and Martinez Arias, 1994; Micchelli, Rulifson and Blair, 1997). Remarkably, Stan localisation was normal (Figure 3H). Therefore, PCP can be established in the absence of secreted Wnt ligands.

### PCP without signals from the wing margin

Our results show that Wnt ligands are not needed for the establishment of the core PCP pathway (Figure 4A). We next determined whether another signal originating from the prospective wing margin could serve as a global cue. This was tested by ablating the wing margin and assaying the effect on PCP (Figure 4B). A combination of Flp, LexA and Gal4-regulated transgenes were used to express Hid and Reaper, two pro-apoptotic proteins (Goyal *et al*., 2000) specifically in the prospective wing margin at third instar stage (See details in Figure 4C-I). Staining with anti-Wingless showed that most of the prospective margin was absent in third instar wing discs (Figure 4C and G). Moreover, the resulting pupal wing completely lacked recognisable margin tissues (Figures 4D and H). The adult wings were particularly small, probably because of lack of Wingless signalling during the growth period. (Figure 4E, F). Yet, PCP, as assayed by Stan staining in pupal wing was normal (Figure 4I), suggesting that no signal emanating from the prospective margin is required for PCP.

**Fig 4.**
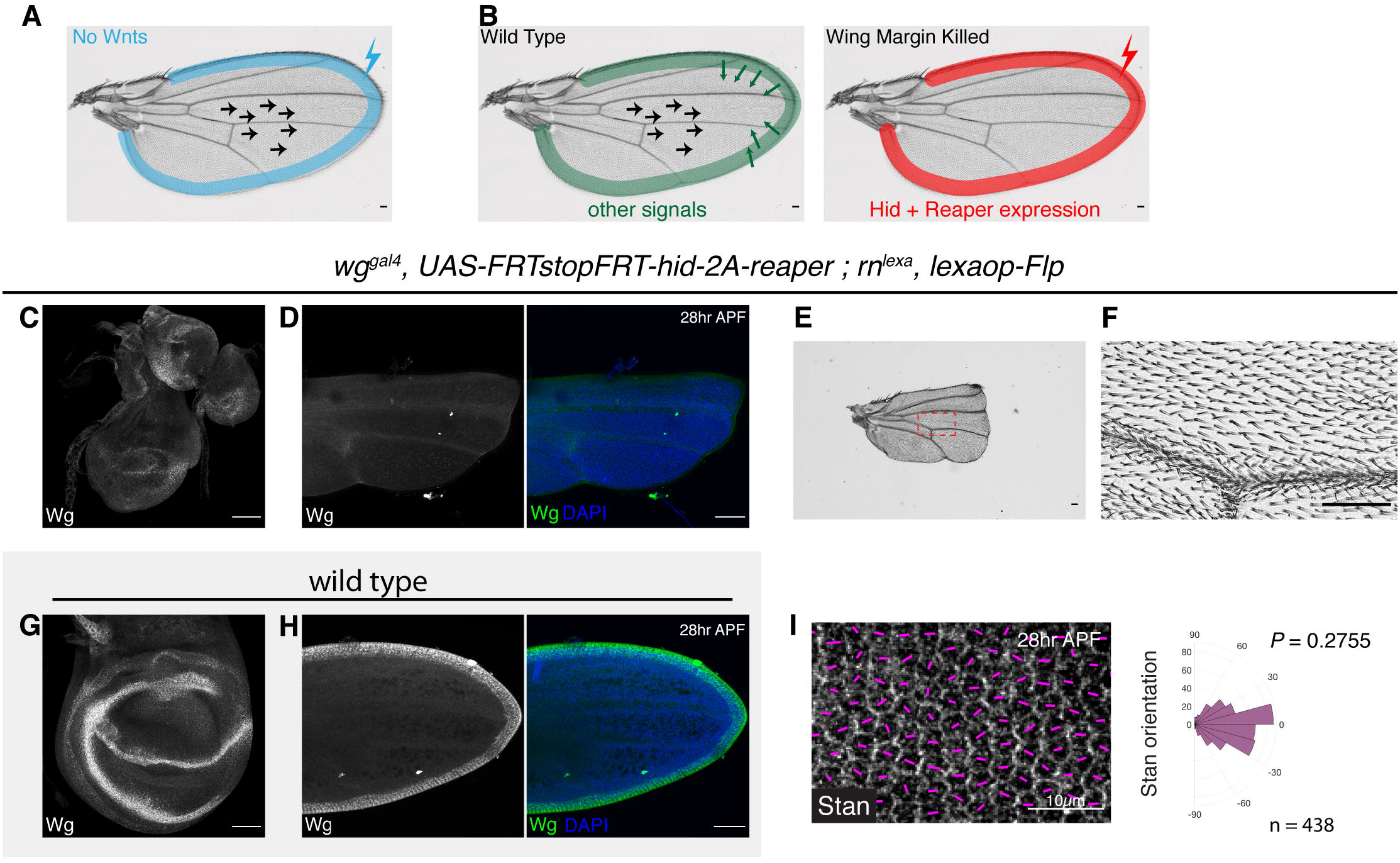
Margin ablation during early third instar does not impair PCP. (**A**) Schematic illustrating that inactivation of all Wnts (expressed in the blue shaded area) is compatible with normal PCP in the wing. (**B**) Experimental set-up to test whether another signal originating from the margin is required for PCP. (**C, D**) Wingless staining in a third instar wing disc and pupal wing with the prospective ablated. Outline of pupal wing is apparent from DAPI staining (blue). (**E, F**) Adult wing with margin ablated. (**G, H**) Wingless immunoactivity in wild type wing disc and pupal wing, for comparison with **C** and **D**. (**I**) Stan polarization in pupal wings with margin ablated.

## Discussion

There has been an ongoing debate whether Wnt ligands play a permissive or instructive role in PCP (Heisenberg *et al*., 2000; Ulrich *et al*., 2005; Witze *et al*., 2008; Gros, Serralbo and Marcelle, 2009; Gao *et al*., 2011; Wu *et al*., 2013; Chu and Sokol, 2016; Minegishi *et al*., 2017; Gao *et al*., 2018; Navajas Acedo *et al*., 2019). To assess rigorously the role of Wnt gradients, we engineered Drosophila larvae so that their wing primordia rely on membrane tethered Wingless as their only source of Wnt. This ensured that any diffusion-based Wnt gradient be eliminated, although at the outset, a gradient based on cytonemes could not be excluded (Stanganello and Scholpp, 2016). To our surprise, wings entirely lacking diffusible Wnt had normal PCP, suggesting that a gradient of Wnt ligand is not needed for Fz-dependent PCP, even if localised ectopic Wnt can orient PCP both in Drosophila wing primordia (Wu *et al*., 2013) and in vertebrate tissues (Chu and Sokol, 2016; Minegishi *et al*., 2017). We suggest therefore that while graded Wnt ligand can orient PCP in these tissues, a Wnt gradient is not necessary. It is possible that ectopic Wnt can hijack the core PCP pathway in a non-physiological manner. Alternatively, a Wnt gradient could normally contribute to PCP but in redundant manner with another global cue. Accordingly, in tissues where no such redundant system exists, a Wnt gradient might be essential, as suggested for the developing mouse limb (Gao *et al*., 2011). In any case, our results suggest that the global alignment of PCP could be achieved without a Wnt gradient.

Despite doubts about the instructive value of Wnt gradients in PCP, it has been generally accepted that, in vertebrates, Wnts are needed, at least in a permissive manner. Yet, as we have shown, in the wing primordia of Drosophila, PCP is established normally in the complete absence of secreted Wnt ligands. This conclusion is based on two sets of experimental results. In one set, Wnt activity was prevented by trapping Wls, and hence all Wnts, in the ER. The effectiveness of this approach can be inferred for the observation that induction of trapping in early primordia completely prevented wing development. Using this approach to trap all Wnts at a later time, but before the period of PCP establishment, we found that PCP can be established normally in the absence of Wnt ligands. This is at odds with the demonstrated essential roles of Wnt5a and Wnt11 in the vertebrate mesoderm. Perhaps, these Wnt ligands have evolved a PCP role that is not present in Drosophila. Our conclusion that Wnt ligands are dispensable for PCP in the Drosophila wing is further strengthened by our observation that PCP is unaffected by complete ablation of the prospective margin, where all the relevant Wnts are produced. The result of our margin ablation experiment also show that PCP can be established without another diffusible cue originating from the prospective margin.

If Wnts are not required for PCP, another global cue must exist. This cannot be from an entirely separate redundant system since removal of Fz1 on its own leads to strong PCP phenotypes. Therefore, any alternative global cue must feed into the Fz-dependent core pathway. Early studies suggested that PCP in the wing relies on graded Fz activity (Vinson and Adler, 1987; Adler, Krasnow and Liu, 1997). Later on, it was proposed that in the *Drosophila* abdomen, each cell is able to compare the proportion of homotypic Stan-Stan and heterotypic Fz-Stan intercellular bridges and that this is transduced into PCP (Lawrence, Casal and Struhl, 2004; Struhl, Casal and Lawrence, 2012). The question remains: how is graded Fz activity established without a gradient of Wnt ligand? A possible role of the Ft/Ds pathway has been suggested several years ago (Adler, Charlton and Liu, 1998; Ma *et al*., 2003; Matakatsu and Blair, 2004). In the wing, the Ft/Ds and Fz core pathways appear to be coupled in a manner that depends on Prickle-Spiny-Legs (Ayukawa *et al*., 2014; Merkel *et al*., 2014; Ambegaonkar and Irvine, 2015), although this may not apply to all tissues (Casal, Lawrence and Struhl, 2006). Thus, a gradient of Ft/Ds could bias Fz polarity especially near the proximal end of the wing where this gradient is most pronounced. The Fz bias could then possibly propagate towards the distal region through short range interaction and/or with the help of tissue flows (Aigouy *et al*., 2010; Aw *et al*., 2016). In the absence of Ft/Ds, PCP is mostly impaired proximally, with distal tissue (near the wing margin) remaining relatively normal. Therefore, it is unlikely that a Ft/Ds gradient is the only global source of PCP orientation. Rather, it is likely that, in wild type wings, the Fz activity gradient integrates proximal Ft/Ds input with distal information from either Wnt ligands or an as yet undetermined global cue. In the wing at least, the Ft/Ds pathway constitutes the major input since normal PCP is established in the absence of the distal signal. Overall, our findings suggest that PCP makes use of redundant global cues, thus ensuring robustness.

## Acknowledgements

We thank Emma Powell, Iris Salecker and Fillip Port for providing fly lines and Joachim Kurth for Drosophila injections. This work was supported by core funding to the Francis Crick Institute (FC001204 to JPV), the Centre National de la recherche Scientifique (CNRS) to YB, the European Union (ERC Advanced TiMorph 340784 to YB) and ARC (SL220130607097 to YB). J.S.Y is supported by the Francis Crick PhD Training Programme. A.M is supported by Fondation ARC pour la recherche sur le cancer (PDF20180507510).

## Experimental Model and Subject Details

### Drosophila strains and fly genetics

Fly strains were raised on standard agar media at 25°C, unless stated otherwise. Strains used in this paper were summarized in the Key Resources Table. DNA injection was performed by either BestGene or the Crick fly facility.

The Wnt reporter toolbox was generated for this study: nlsGFP reporters for DWnt2, DWnt4, DWnt6, and DWnt5; *DWnt10[HA]*; *DWnt10[GAL4]*; *DWntD[GAL4]*. Other fly lines generated in this study include: *DWnt4[KO], wg[cNRT]*; *DWnt2[KO], DWnt4[cKO], wg[cNRT], DWnt6-KO], DWnt10[KO]*; *fz1^P21^*; *UAS-Nanobody^KDEL^*; *wls[ExGFP]*; *dsh[cKO]*; rn^lexa^. *wls^sgRNA^* and *UAS-Cas9* were gifts from Fillip Port (Port *et al*., 2020). *UAS-FRT stop FRT-hid-2A-reaper*, *lexaop-Flp* and *UAS-Flp5* were gifts from Iris Salecker. *wg^gal4^* used in this study was as described in (Alexandre, Baena-Lopez and Vincent, 2014). *wg::GFP* was a gift from Simon Bullock (Port *et al*., 2014). The following stocks were obtained from the Bloomington Drosophila Stock Centre: *rn^gal4^*; *UAS-Flp*; *nub^gal4^*; *UAS-GFP*.

### Genotypes

Figure 1

(B) *w1118*
(C) *wg[cNRT]; rn^gal4^, UAS-Flp*
(D) *DWnt4[KO], wg[cNRT]; rn^gal4^, UAS-Flp*
(E) *fz1^P21^*

Figure 2

(A-B) *wg::GFP*
*nlsGFP-PQR-DWnt* (for DWnt2, DWnt4, DWnt5, and DWnt6)
*DWnt10[HA]*
(C-D) *DWnt2[KO], DWnt4[cKO], wg[cNRT], DWnt6-KO], DWnt10[KO]; rn^gal4^, UAS-Flp, UAS-Flp5*

Figure 3

(B) *vg^gal4^, UAS-Nanobody^KDEL^*
(C) *wls[ExGFP]*
(D) *GFP-wg*
(E-H) *wls^sgRNA^, nub^gal4^, UAS-Cas9; wls[ExGFP], UAS-Nanobody^KDEL^*

Figure 4

(C-F, I) *wg^gal4^, UAS-FRT stop FRT-hid-2A-reaper; rn^laxa^, lexaop-Flp*
(G) *w1118*

Supp Figure 1

(E) *DWnt10[GAL4]/+; UAS-GFP/+*
(F) *DWntD[GAL4]/UAS-GFP*

Sup Figure 3

(D-E) *dsh[cKO]/Y; nub^gal4^, UAS-Flp*

## Method Details

### Generation of DWnt4[KO] in wg[cNRT] background

The *wg[cNRT]* conditional allele (*FRT wg FRT nrt-wg*) was generated by replacing the endogenous *wg* locus with *FRT wg FRT nrt-wg*, as described in (Alexandre, Baena-Lopez and Vincent, 2014). The *DWnt4[KO], wg[cNRT]* chromosome was generated via CRISPR-Cas9 and homologous recombination mediated repair in the *wg/cNR’/’/*background. The first exon of *DWnt4* was replaced with *attP* and *pax-GFP*, a selection marker, using the pTV^*GFP*^ targeting vector with 1.2kb 5’ and 1.5kb 3’ homology arms (Figure S1A). CRISPR target sites were chosen in unconserved regions, one upstream of the 5’UTR (ATGAGCAAAATGCAATCTAT), one in the intronic region following exon 1 (AGCATTTGAGGACGGCAAAC). Primers used for 5’arm: CATTAGAATTCAGCGCGTCTAAATGGACAC (Forward); TAATGGGTACCCTCAAATGCTATTATAATTCTGAAAAGTTTATAATAATC (Reverse). Primers used for 3’arm: CATTAACTAGTGCAATCTATAGGTGACTTTAACAATCAG (Forward); TAATGACGCGTTGTGTATGTTTTCTCCTCCG (Reverse). The resulting pTV^*GFP*^-DWnt4[KO] construct and the *DWn4^sgRNA^* donor vector were co-injected into embryos from a cross of *nanos-Cas9* line and *wg[cNRT]*. Successful transformants were identified by GFP expression in the eyes, and subsequently PCR verified.

### Generation of DWnt2, DWnt4, DWnt6 and DWnt5 GFP reporters

To generate the nls-GFP reporters of DWnt2, DWnt4, DWnt6 and DWnt5 expression, a nls-GFP-T2A targeting vector was built by introducing an nls encoding sequence (CCTAAGAAGAAGCGGAAAGTA) and a T2A sequence (GAGGGCCGCGGCTCCCTGCTGACCTGCGGCGACGTGGAGGAGAACCCCGGCCCC) upstream and downstream of GFP sequence, respectively in the CHE929^*GFP-lox-mini-white-lox*^ vector, which allows transformant selection with *mini-white* (Pinheiro *et al*., 2017). At least 1kb of 5’ and 3’ homology arms were cloned using the following primers:

DWnt2 5’arm:
CCCGGGCTAATTATGGGGTGTCGCCCTTCGCTCAGTTCTTATACGACACTCGCACC
(Forward); GCTCACTACTTTCCGCTTCTTCTTAGGCATCGAGCCCAGCAGCTGGCACTAT
(Reverse)
DWnt2 3’arm:
TGCGGCGACGTGGAGGAGAACCCCGGCCCCATGTGGAAAATACATAACAAGCTCTTAAT
C (Forward);
GCCCTTGAACTCGATTGACGCTCTTCGGGGAGTTACAAAACTCGTTTAATAATATGTATC
(Reverse)

DWnt4 5’arm:
CCCGGGCTAATTATGGGGTGTCGCCCTTCGTGGGAAAGCGTGAAAACTTCTGGCATC
(Forward);
GCTCACTACTTTCCGCTTCTTCTTAGGCATGAAGCGGTTGTTGGTGAATTGTGGATT
(Reverse)
DWnt4 3’arm:
TGCGGCGACGTGGAGGAGAACCCCGGCCCCATGCCATCGCCAACGGGAGTCTTT
(Forward) and
GCCCTTGAACTCGATTGACGCTCTTCGGGGGATCTGACTTGTTCTAGCCATAAATTGTCG
(Reverse)

DWnt5 5’arm:
CCCGGGCTAATTATGGGGTGTCGCCCTTCGGGTATCTATTATTCGAGGTGGATTAGTG
(Forward);
AACAATACTCTGTAATAGTAATAGTAAGAGATGCCTAAGAAGAAGCGGAAAGTAGTGAG
C (Reverse)
DWnt5 3’arm:
TGCGGCGACGTGGAGGAGAACCCCGGCCCCATGAGTTGCTACAGAAAAAGGCACTTTCT
(Forward);
GCCCTTGAACTCGATTGACGCTCTTCGGGGGTGTCTCGTTGAGCAGTATCACCTG
(Reverse)

DWnt6 5’arm:
CCCGGGCTAATTATGGGGTGTCGCCCTTCGCAGATCCATATGAACGGTCAGCGC
(Forward); GCTCACTACTTTCCGCTTCTTCTTAGGCATAGCCGCCAAATGCAGCGGCAACAA
(Reverse)
DWnt6 3’arm:
TGCGGCGACGTGGAGGAGAACCCCGGCCCCATGCGTTTGCTCATGGTAATTGCAATTTTA
(Forward);
GCCCTTGAACTCGATTGACGCTCTTCGGGGCCTCCGCGGTTTGTTTATTTATTTATTCAT
(Reverse)

Target sites were chosen using http://targetfinder.flycrispr.neuro.brown.edu/ website (Gratz *et al*., 2014) to avoid off-target sides. sgRNAs were cloned in *pCFD5: U6:3-t::gRNA* vector (Port *et al*., 2014). For all the reporters, the sgRNA vector and the *nls-GFP-T2A* targeting vector were injected into vas-Cas9 lines (Gratz *et al*., 2014) by Bestgene, and subsequently PCR verified.

### Generation of DWnt10[HA] and GAL4 lines

The *DWnt10[HA]* allele was generated via two rounds of modifications (Figure S1C). In the first step, the first exon of *DWnt10* was replaced with *attP* and *pax-Cherry*, using the pTV^*Cherry*^ targeting vector with 1kb 5’ and 1.5kb 3’ homology arms. Target sites were chosen in unconserved regions, one upstream of the 5’UTR (TGCTTTAAATACAAGAATGC), one in the intronic region following exon 1 (TGAGATAAGAAGATGTTCAG). Primers used for 5’arm: CATTAGAATTCAGCTCGGCTCTCGTCTCATAG (Forward);

TAATGGGTACCAAAGCATGGAAATTAAGGGAAATATAC (Reverse). Primers used for 3’arm: CATTAGACGTCAGAAGATGTTCAGAGGCAGAC (Forward);

TAATGACCGGTTAACTAAAACTAAGGCGATGTACTCATG (Reverse). The resulting pTV*^Cherry^-DWnt10[KO]^attP^* and *DWnt10^sgRNA^* vectors were co-injected into embryos from the *nanos-Cas9* line. Successful candidates were identified by Cherry expression in the eyes and subsequently PCR verified. This created a null allele of *DWnt10* (*DWnt10[KO]^attP^*). The attP site was then used for reintegration of RIV^*white*^ (Baena-Lopez *et al*., 2013) modified as follows. A DNA fragment containing the 5’UTR, CDS, and 3’UTR of DWnt10 was synthesized by GeneWiz, with the sequence of HA-tag inserted in an unconserved region in exon 6. This fragment was cloned into the RIV^*white*^ vector with the primers CATTAGAATTCTCATAGTCGCGCGACTCG (Forward) and TAATGGGTACCGACAGACTAATTATGCATCTTATTGTTCGTATTTACAAAATATATTT (Reverse). The RIV^*white*^-DWnt10-HA vector was then injected into the *DWnt10[KO]^attP^* line to generate *DWnt10[HA]* via PhiC31-mediated integration.

*DWnt10[GAL4]* was generated using the same strategy as for *DWnt10[HA]*, but instead of the RIV^*white*^ integration vector, a RIV^*gal4*^ integration vector (Baena-Lopez *et al*., 2013) was inserted into the *attP* site of *DWnt10[KO]^attP^* (Figure S1D).

*DWntD[GAL4]* was generated via two rounds of modification (Figure S1D). First, exon 1 was replaced with *attP* and *pax-Cherry*, using the pTV^*Cherry*^ targeting vector with 1.5kb 5’ and 1.5kb 3’ homology arms. *DWntD^sgRNA^* was designed to target sites in unconserved regions, one upstream of the 5’UTR (GCTATATAAGTGTGCTGACC), one downstream of the 3’UTR (GTTTTAGCTACAGGTGGTTT). Primers used for amplification of the 5’ homology arms were: CATTAGCGGCCGCATGTAGTGGGGGAATCTCAAG (Forward); TAATGGGTACCACACTTATATAGCCTGCAAATCCC (Reverse).

Primers used for 3’homology arm: CATTAGACGTCCTACAGGTGGTTTAATAATTACGGATTTTAGAGT (Forward); TAATGACGCGTCTGATGCAGCGGCCAC (Reverse). The pTV*^Cherry^-DWntD[KO]^attP^* and *DWntD^sgRNA^* plasmids were co-injected into embryos from the *nanos-Cas9* line to generate the null allele *DWntD[KO]^attP^* (PCR verified). A RIV^*gal4*^ integration vector (Baena-Lopez *et al*., 2013) was then introduced by PhiC31-mediated integration to generate *DWntD[GAL4]* (Figure S1D).

### Generation of multiple DWnt mutants in wg[cNRT] background

First, a double knockout of *DWnt6 and DWnt10* was sequentially generated on the *wg[cNRT]* chromosome by injecting sgRNAs targeting the first exon of both genes in *wg[cNRT] nos-Cas9 embryos*. Details of the target sites and cloning strategies are described in Figure S2B. Indels were screened by genomic DNA extraction and PCR sequencing of homozygous candidates.

Next, a conditional *DWnt4* allele, *DWnt4[cKO]*, was generated by CRISPR-Cas9 and homologous recombination mediated repair on the *wg[cNRT], DWnt6[KO], DWnt10[KO]* chromosome. DWnt4 was made conditional by replacing the 5’UTR and exon 1 with the same sequence but flanked by FRT71 sites (Figure S2B). CRISPR target sites chosen in unconserved regions, one upstream of the 5’UTR (ATGAGCAAAATGCAATCTAT), one in the intronic region following exon 1 (AGCATTTGAGGACGGCAAAC). The rescuing pTV^*GFP*^-DWnt4[cKO] construct was made by first generating PCR fragments encoding the 5’ arm and exon 1 of DWnt4. They were then stitched together and inserted into pTV^*GFP*^ upstream of the pax-GFP selection cassette by Gibson Assembly. FRT71 sites were included in the primers so that they would be inserted between the 5’arm and the rescuing exon 1 and also immediately after the rescuing exon 1. The 3’ arm was then amplified by PCR and inserted after the pax-GFP selection cassette. Primers for 5’ arm: GCGAATTGGAGCTGAATTCCTGTAGATAAACCCTTGGGG (Forward); TCTAGAAAGTATAGAAACTTCAGGTGACTTTAACAATCAGGTGATTG (Reverse). Primers for DWnt4 exon 1 region flanked by the homology arms: GAAGTTTCTATACTTTCTAGAGAATAGAAACTTCATCGATTGCATTTTGCTCCTAATATCTTCAATTTCCACACATTTTTCATTTGAGA (Forward);

GTTGGGGCACTACGGGTACCGAAGTTTCTATTCTCTAGAAAGTATAGAAACTTCAGCATT TGAGGACGGCTAACATGAAA

TCAATATTTTCGATTTCTAGAATTTTGCC (Reverse). Primers for 3’arm: CATTAACTAGTATTATAATTCTGAAAAGTTTATAATAATCGCTTCCTG (Forward); TAATGACGCGTGCTTCAAGCTTATACGCACAGAA (Reverse). The resulting pTV*^GFP^-DWnt4[cKO]* plasmid was co-injected with *DWn4^sgRNA^* into embryos from a cross of *nanos-Cas9* line and the *wg[cNRT], DWnt6[KO], DWnt10[KO]* line. Successful candidates were identified by pax-GFP expression and subsequently PCR verified.

A DWnt2[KO] was generated separately by replacing the first exon with *attP* and *pax-Cherry*, using the pTV^*Cherry*^ targeting vector with 1.5kb 5’ and 1.5kb 3’ homology arms (Figure S2B). CRISPR target sites were chosen in the unconserved regions, one upstream of the 5’UTR (AGTAGTAGTACTACTTGATC), one in the intronic region following exon 1 (AAATCAAAATACCTTCATCG). Primers used for 5’arm: CATTAGCGGCCGCATTATCTGGAATTTAAAAGTCTTGGCACAC (Forward); TAATGGGTACCGTACTACTACTATCCAACTTTCCACTCT (Reverse). Primers used for 3’arm: CATTAACGCGTGTATTTTGATTTTGGTGAAATGCTTGAAAAG (Forward); TAATGACCGGTCATAAAGGCTGTTCAACATGCC (Reverse). The resulting pTV*^Cherry^-DWnt2[KO]^attP^* plasmid was co-injected with *DWnt2^sgRNA^* into *nanos-Cas9* embryos. Successful candidates were identified by pax-Cherry expression and subsequently PCR verified. *DWnt2[KO]* was then recombined with *DWnt4[cKO], wg[cNRT], DWnt6[KO], DWnt10[KO]*. Successful recombination was screened by the presence of pax-GFP (from *DWnt4[cKO]*), and extra bright pax-Cherry signal in the eye, as both the *DWnt2[KO]* and *wg[cNRT]* alleles harbour the pax-Cherry marker. The recombinant was subsequently verified via PCR.

### Generation of dsh[cKO]

The *dsh[cKO]* allele was generated in two steps (Figure S3B). First, *pTV^CCìerry^* with a 2kb 5’arm and 1kb 3’arm was used to replace the coding region with an attP site and pax-Cherry to generate *dsh[KO]^attp^*. CRISPR target sites were chosen in the 5 ‘UTR and just after the stop codon ((TTCCCGTGGATTTCCGCAGT, CGCAGTCGGCGCAGCTAAAA, CTACAATACGTAATTAAATA and TACGGATACGTCCTGATCGT). Primers used for 5’arm: ATGCAGGCGGCCGCTATCAGCCGTCGTGCGTG (Forward);

TGACTCACATATGCCTCGACGCGATAAAACGCGATC (Reverse). Primers used for 3’arm: TGCACCTAGGTGAGATTGGTTTCTTTAACGCATTTTAGCTG (Forward);

CGTCAACCGGTCATTTCGTATTAGTTGTATGTACGGAAGTTGAC (Reverse). The resulting pTV*^Cherry^-dsh[KO]^attp^* plasmid was co-injected with *dsh^sgRNA^* into *nanos-Cas9* embryos. Successful candidates were identified by pax-Cherry expression and subsequently PCR verified. This created a null allele of *dsh* (*dsh[KO]^attp^*). The attP site was then used for reintegration of *dsh-GFp* flanked by FRT sites using RIV10^*dsh-GFp*^.

RIV10^*dsh-GPP*^ generated by first cloning *dsh* from genomic DNA into *pBS-KS*. Primers used were; ATGGTCAGCTAGCGATCGCGTTTTATCGCGTCGAGGAGTTTTCCCGTGGATTTCCGCAGTCGGCGCAGCTAAA (Forward) and TCCATGACCGGTCCATGCCCGGAATGTTTCCG (Reverse). The pBS-KS^*dsh*^ vector was then opened using a unique SnaBI site just prior to the stop codon of dsh. GFP was amplified via PCR from pEGFP-N1 (Clontech) and inserted into the linearized pBS-KS^*dsh*^ vector using Gibson Assembly to generate pBS-KS^*dsh-GFP*^. Primers for cloning GFP into pBS-KS^*dsh*^:

CGTATCCGTATTTAATTACGTATTGGGAGGTTCGGGAGGTGGTTCGGGAGGTGGAGGTTCGGGAGTGAGCAAGGGCGAGGA (Forward); GAAAGGTTTCTACAATACgtaCTTGTACAGCTCGTCCATGCC (Reverse).

Dsh contains a ‘YV?’ PDZ motif at the carboxy-terminus that has been suggested to be essential for function (Lee, Shi and Zheng, 2015). This ‘YV? motif was hence duplicated and inserted after the GFP coding sequence. The *dsh-GFP* sequence was then subcloned from the pBS-KS^*dsh-GFP*^ vector into RIV10^*attB-GFP*^ via the Nhel and Agel restriction sites to generate RIV10^*dsh-GFP*^ The RIV10^*dsh-GFP*^ vector was then injected into the *dsh[KO]^attP^* line to generate the *dsh[cKO]* line via PhiC31-mediated integration.

### Generation of UAS-Nanobody^KDEL^

*UAS-Nanobody^KDEL^* was generated by PCR amplifying the coding region of the VHH4 nanobody from *pHT201* (gift from Dr. Peter Thorpe, Queen Mary, University of London), and subcloning it into *pUAST*. The sequence encoding <DEL was contained within the reverse primer used for amplification of the nanobody, such that the <DEL was located at the C-terminal end of the nanobody just prior to the stop codon. pUAST-*Nanobody^KDEL^* was then randomly integrated via P-element insertion and one line on the third chromosome was recovered.

### Genersation of wls[ExGFP]

The *wls[ExGFP]* line was generated by two rounds of injection (Figure S3F). First, using the accelerated ‘Ends out’ homologous recombination method described in (Baena-Lopez *et al*., 2013), a region comprising the three exons of *wls* (from 20bp upstream of the initiation codon till 49bp after the stop codon) was replaced by an attP site and pax-Cherry, using the *pTV^Cherry^* integration vector. In a second step, a DNA fragment encoding *wls[ExGFP]* was generated starting with Lit28-EVI^2XHA^, which was generated as follows. DNA encoding Evi2HA was synthesized by Genewiz, such that two HA tags flanked by Gly/Ala linker were inserted between amino acid 506(D) and 507(N) s. The native 5’UTR and the 3’UTR of *wls* were subsequently added to generate an evi2HA cDNA. The resulting Lit28-EVI^2XHA^ was digested with AatII. This allowed the HA tags to be replaced with GFP (amplified with primers including the AatII sites from pEGFP-N1 (Clontech). Forward primer: TGACGACGTCGTGAGCAAGGGCGAGGAGCTGTTCA; reverse primer: GTCGACGTCCTTGTACAGCTCGTCCATGCCGAG. DNA encoding Wls[ExGFP] was then subcloned from Lit28 into RIV^*Cherry*^, which was subsequently injected into the *wlsKO[*attP] to generate the *wls[ExGFP]* line via PhiC31-mediated integration.

### Generation of rn^lexa^

Rn-LexA was generated using RMCE (Venken *et al*., 2011), to insert *pBS-KS-attB2-SA-T2A-LexA::GADfluw-Hsp70* into a *rn* MIMIC line (Bloomington, #44158). *pBS-KS-attB2-SA-T2A-LexA::GADfluw-Hsp70* was a gift from Benjamin White (Addgene plasmid # 78304). (Diao *et al*., 2015)

### Immunostaining and image acquisition

The primary antibodies used were: mouse anti-Stan (1:10, pre-adsorbed, DSHB Flamingo #74), rat anti-Shg (1:50, pre-adsorbed, DSHB DCAD2); mouse anti-Wg (1:500, DSHB 4D4), rabbit anti-GFP (1:500, Abcam ab6556). Secondary antibodies (Alexa Fluor 488, 555, 647) were used at 1:200.

Larval wing discs and pupal wings were dissected and fixed in PBS 4% formaldehyde for 20min (larval discs) or 1hr (pupal wings). Dissected tissues were then washed in 0.1% PBT three times and then incubated in blocking solution (0.1% BSA) for 1hr. Samples were incubated in primary antibodies overnight at 4°C, and then washed in PBT three times. Secondary antibodies in blocking solution were then added and incubated for 2hr at room temperature. Samples were then washed three times in PBT, and then mounted in Vectashield with DAPI. All immunofluorescence images were acquired from a Leica SP5 confocal microscope. Adult wings images were obtained from a wide field microscope (Zeiss Axiovert 200M).

### Quantification and statistical analysis

#### Polarity measurement

Z-stacks of the acquired images of each wing was projected using a MATLAB script, modified from (Heller *et al*., 2016), which allowed the projection of the apical regions of epithelial tissues into 2D images, based on the anti-Ecad immunofluorescent staining. Membrane masks were generated using Tissue Analyzer (Aigouy *et al*., 2010). To determine the asymmetric localisation of Fmi of individual cells, a MATLAB script was used (Strutt, Gamage and Strutt, 2016), based on Fmi staining intensity in the pupal wing. This script also generated PCP nematics for each individual cell, which takes into account the orientation and magnitude of the Stan polarization. This nematic order was visualised as magenta lines superimposed onto the original anti-Stan immunofluorescence image. Polar histograms were generated in MATLAB to visualise the orientation of Stan, such that 0° was oriented as pointing distally in the pupal and adult wings.

#### Statistical analysis

The two sample Kolmogorov-Smirnov test was used as a statistical test to compare the difference in the distributions of cellular polarity of cells from two independent samples that contain nonindependent data, such as the case in PCP analyses when comparing wild type versus mutant conditions.

## Notes

### Competing Interest Statement

The authors have declared no competing interest.

