## Supplementary_Figures for "Frizzled-dependent Planar Cell Polarity without Wnt Ligands"

***Supp Fig. 1****: Reporters of DWnt expression (related to Figure 1D and 2A-B)*

(**A**) Generation of *DWnt4* mutant by removing the first exon (in the *wg[cNRT]* background). CRISPR target sites (vertical black bars) were chosen in unconserved regions flanking the 5'UTR and first exon. Details for cloning strategies of all transgenic lines generated in this paper are described in the Methods section. (**B**) Generation of nlsGFP-expressing reporters for DWnt2, DWnt4, DWnt5, and DWnt6. (**C**) Generation of *DWnt10[HA]*. (**D**) Generation of Gal4 lines for *DWnt10* and *DWntD*. (**E, F**) Expression patterns of *DWnt10[GAL4]* and *DWntD[GAL4]*, as reporter with *UAS-GFP* expression. Scale bars are 50μm.

***Supp Fig 2:*** *A strain for the manipulation of five DWnt genes simultaneously (related to main Figure 2C-D)*

(**A**) Schematic diagram of the generation of a quadruple DWnt mutants in the *wg[cNRT]* background. The *wg[cNRT]* allele was marked with *pax-Cherry*, which allowed us to exclude multi-genic deletions. The conditional DWnt4 mutant was marked with *pax-GFP*, another selection marker. (**B**) Details of the mutations generated in each *DWnt*. INDELS were created in *DWnt6* and *DWnt10*; the first exon of *DWnt2* was removed; the first exon of DWnt4 was flanked by FRT71 sites, which allowed the tissue specific Flp-excision of the region to create a *DWnt4* mutant.

**Supp Fig. 3:** *Growth and PCP activities of Dsh can be temporally separated (related to main Figure 3)*

(**A**) Comparison of the canonical Wnt pathway and the Fz/PCP pathway. Fz and Dsh are common components of both pathways. (**B**) Schematic of the generation of the Dsh conditional null allele (*dsh[cKO]*). (**C**) A timeline of Drosophila development showing the approximate periods when growth occurs, and PCP is specified. The time of activation of relevant wing drivers is also shown. (**D, E**) Adult wing and pupal wing Stan polarization upon inactivation of *dsh[cKO] with* $nub^{gal4}$ and *UAS-Flp*. (**F**) Generation of *wls[ExGFP]*, with GFP inserted into the fourth extracellular loop of Wls.

**Supplementary Figure 1.** Reporters of DWnt expression (related to main Figure 1D and 2A-B)

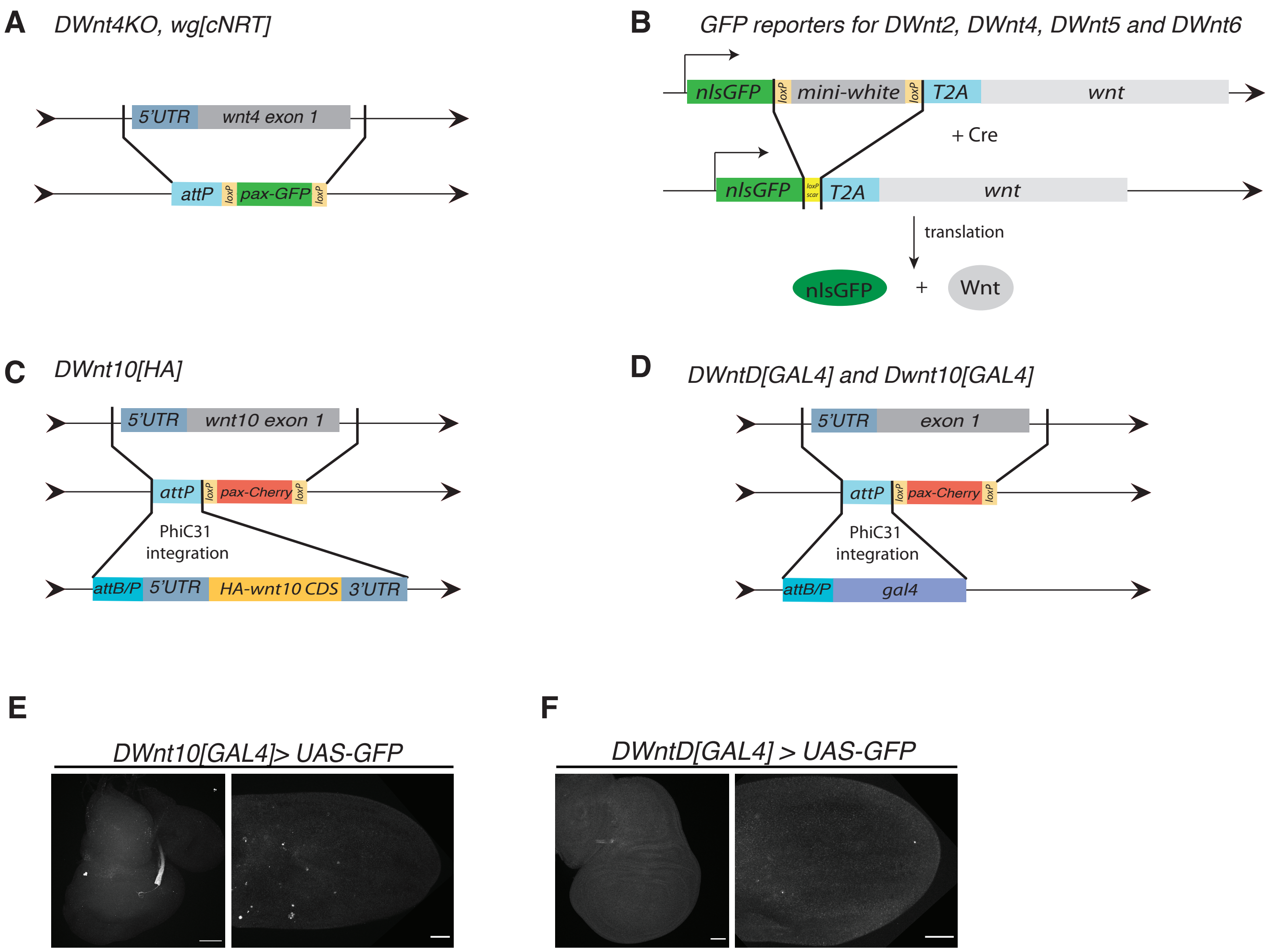

**Supplementary Figure 2**. A strain for the manipulation of five DWnt genes simunltaneously (related to main Figure 2C-D)

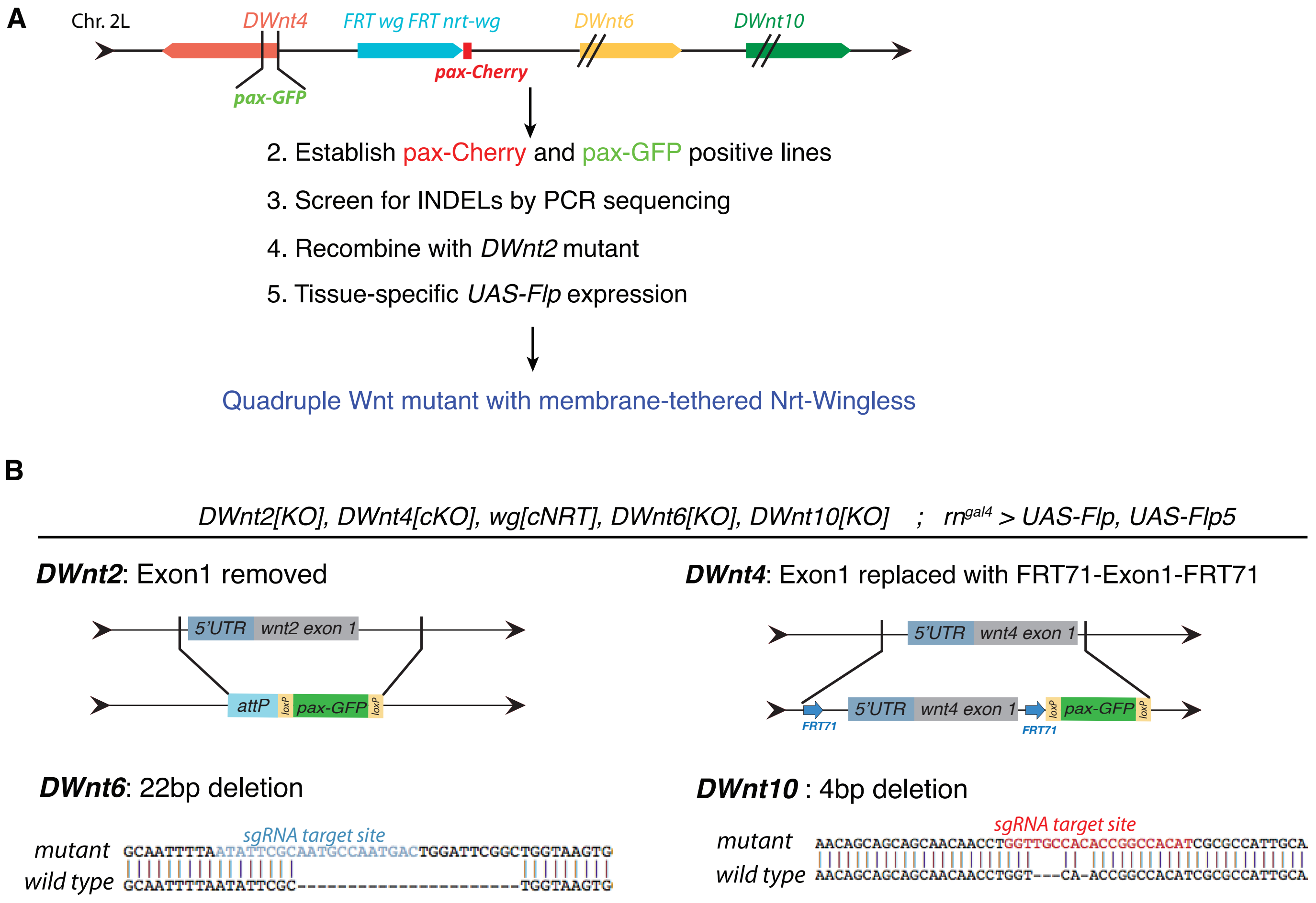

**Supplementary Figure 3.** Growth and PCP activities of Dsh can be temporally separated (related to main Figure 3)

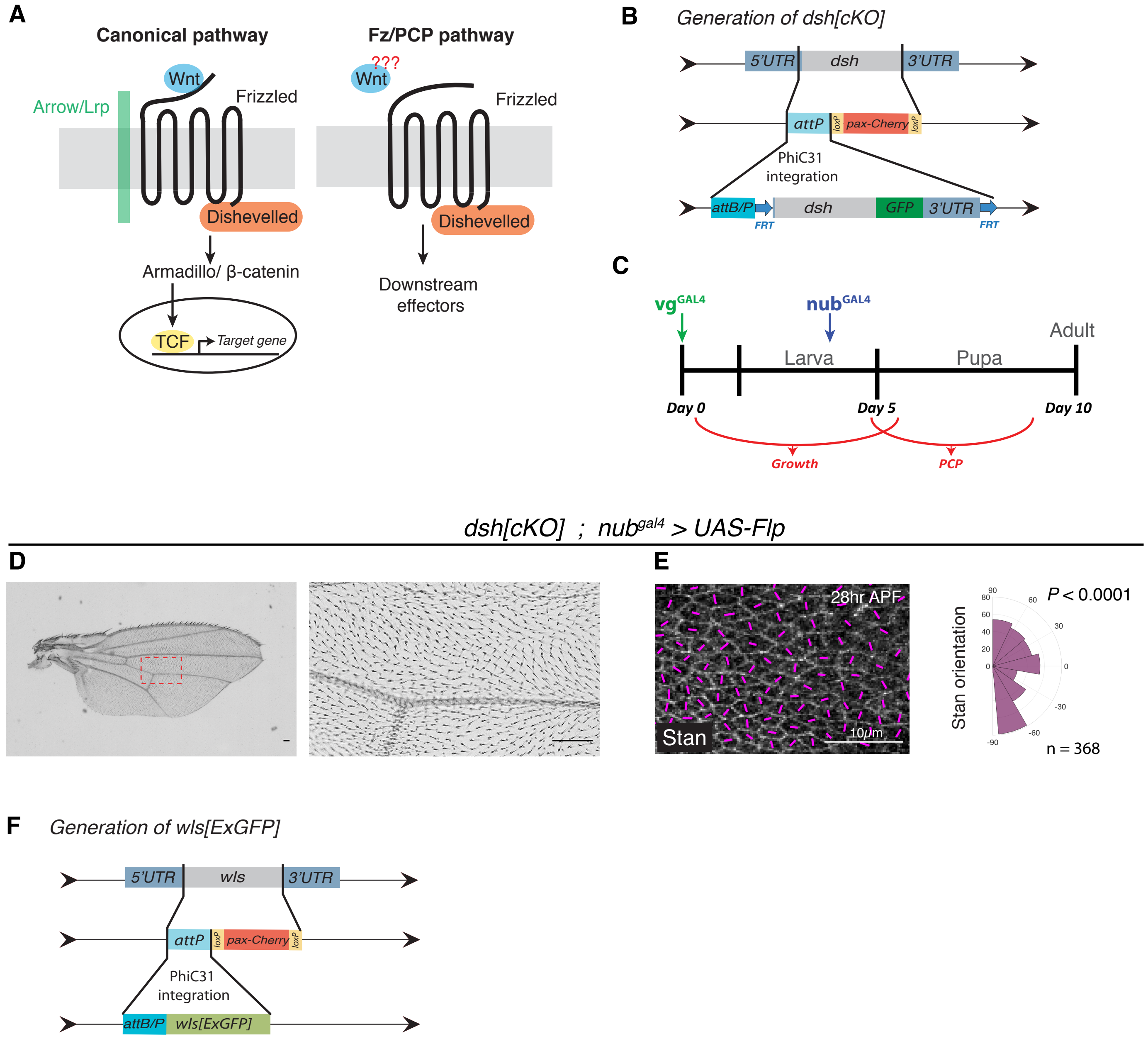
